# Self-organization of high-dimensional geometry of neural activity in culture

**DOI:** 10.1101/2025.01.14.630383

**Authors:** Asahi Nakamuta, Dai Akita, He Zhang, Yuta Kawahara, Hirokazu Takahashi, Jun-nosuke Teramae

**Affiliations:** Graduate School of Informatics, Kyoto University, Japan; Graduate School of Information Science and Technology, University of Tokyo, Japan

## Abstract

A vast number of neurons exhibit high-dimensional coordination for brain computation, both in processing sensory input and in generating spontaneous activity without external stimuli. Recent advancements in large-scale recordings have revealed that this high-dimensional population activity exhibits a scale-free structure, characterized by power law and distinct spatial patterns in principal components (PCs). However, the mechanisms underlying the formation of this high-dimensional neural coordination remain poorly understood. Specifically, it is unclear whether the characteristic high-dimensional structure of population activity emerges through self-organization or is shaped by the learning of sensory stimuli in animals. To address this question and clearly differentiate between these two possibilities, we investigated large-scale neural activity in dissociated neuronal culture using high-density multi-electrode arrays. Our findings demonstrate that the high-dimensional structure of neural activity self-organizes during network development in the absence of explicit sensory stimuli provided to animals. As the cultures mature, the PC variance exhibits a power-law decay, and the spatial structures of PCs transition from global to localized patterns, driven by the temporal correlations of neural activity. Furthermore, we observed an unexpected co-occurrence between the power-law decay in PCA and neuronal avalanches, suggesting a link between self-organized criticality and high-dimensional activity. To uncover the mechanism behind this co-occurrence, we developed a new theoretical framework and demonstrated that it arises from heavy-tailed synaptic connectivity. By highlighting a developmental origin of the high-dimensional structure of neural activity, these findings deepen our understanding of how coordinated neural computations are achieved in the brain.

**Significance:** One of the most intriguing questions in neuroscience is how the extremely high-dimensional yet efficiently orchestrated coordination of neural activity is organized. Central to this question lies the “nature versus nurture” debate: Is the high-dimensional coordination self-organized during development, or is it acquired through learning in animals? We take a novel approach by applying cutting-edge analytical techniques to study high-dimensional neural activity in dissociated neuronal cultures. Our findings reveal that fundamental features characterizing the high-dimensional coordination of neurons, including the scale-free geometry of the neural manifold and the intrinsic spatial structure of neural dynamics, are self-organized without sensory stimuli. Remarkably, we also discover that this high-dimensional geometry emerges concurrently with neuronal avalanches, a hallmark of self-organized criticality in neural circuits.

## 1 Introduction

To perform the computational tasks essential for animal survival, cortical neurons generate coordinated spiking activity rather than responding independently to input signals. Previous studies, which measured the activity of a limited number of neurons, hypothesized that this coordinated neural activity is constrained to low-dimensional subspaces due to strong correlations between neurons [1, 2]. However, recent advances in large-scale recording techniques have revealed that the activity is, in fact, high-dimensional [3, 4, 5, 6, 7, 8]. The number of dimensions required to explain a fraction of the variance in coordinated activity consistently increases with the number of recorded neurons. This intrinsic high-dimensionality of coordinated neural activity was first reported in stimulus-driven responses of neurons in the visual cortex [3]. Subsequent studies have further elucidated the ubiquity of this high-dimensional structure, as it is observed across brain regions and species, even in spontaneously generated neural activity in the absence of sensory stimuli in animals [4, 6, 7, 8, 9].

This high-dimensional population activity exhibits two distinct features observed through principal component analysis (PCA): a power-law decay of explained variance and a characteristic spatial structure of principal components (PCs). Experiments have shown that the decay of the rank-ordered explained variance of PCs is well fitted by a power-law function. This power-law decay indicates that the contribution of PCs to neural activity decreases surprisingly slowly as the rank increases. Therefore, this observation clearly shows that the effective dimensionality of neural activity continues to grow as the number of observed neurons increases [4, 6]. The spatial patterns of neurons that contributed to or participated in each PC vary qualitatively as the rank of the PCs changes. Neurons that participate in lower-ranked (i.e., dominant) PCs are spatially localized within brain regions. In contrast, neurons that participate in higher-ranked (i.e., minor) PCs are globally and sparsely distributed, spanning the entire cortex [7]. This result is particularly intriguing because it is consistent with recent findings suggesting that cognitive processes emerge from multi-scale neural activity, ranging from local regions to the entire brain [10].

While high-dimensional neural activity has been observed across various species and brain regions, its underlying formation mechanism remains poorly understood. Specifically, it is still unclear whether the power law and the spatial structure in PCA are self-organized during development or acquired through experience. Solving this question is crucial for uncovering the origin and function of the observed high dimensionality. Previous in vivo studies have failed to address this due to methodological constraints. Although some studies have investigated the dependence of correlation and spatial extent of zebrafish tectal spontaneous activity on retinal inputs [11, 12], they did not adequately control for the influence of other sensory modalities and lacked a perspective on high-dimensionality. Moreover, a previous study demonstrated that high-dimensional activity is not directly inherited from the input by altering visual stimuli presented to mice [3]; however, it did not examine the contribution of the mice’s prior visual experiences before the experiments.

To address this issue, we utilized a dissociated neuronal culture [13, 14], in which cortical neurons are initially dissociated and then develop into network. This setup allows for the observation of neuronal network dynamics in a simplified, controlled environment, free from the constraints of development and prior learning. Importantly, the dissociated culture preserves key biological features of the cortex, such as the approximate 4:1 ratio of excitatory to inhibitory neurons [15, 16, 17, 18] and the major neurotransmitters (glutamate, GABA, and acetylcholine) [13]. This similarity suggests that the culture captures essential aspects of cortical organization, making it a suitable model for studying intrinsic self-organized network dynamics. A multi-electrode array embedded in the dish [19, 20, 21] allows high-throughput recordings of the activity.

Using this setup, we found that the high-dimensional coordination of spontaneous activity is self-organized during development. As the cultured networks matured, the power law and spatial structure in PCA spontaneously emerged. Furthermore, by observing the process through which the high-dimensionality is established, we discovered that the temporal correlation of neural activity, which arises during the development of the neuronal network, plays a crucial role in the self-organization of the characteristic high-dimensional structure.

We further discovered that the power law in PCA concurrently emerges with neuronal avalanches, a distinct power law in synchronized firing events. While the neuronal avalanches have been extensively studied as a self-organized criticality (SoC) in neural systems [22, 23, 24, 25, 26], their relationship to high-dimensional activity has remained largely unknown, as they have been studied independently. Our results revealed the co-occurrence of the power law in PCA and neuronal avalanches, suggesting a potential link between SoC and the emergence of high-dimensional neural activity. Since these two power laws focus on different aspects of neural activity and we could easily constitute examples where they do not co-occur, their co-occurrence is a remarkable finding. Furthermore, we theoretically investigated the underlying mechanism of this co-occurrence. Using mathematical modeling and analytical calculations, we revealed that heavy-tailed connectivity, widely observed in vivo [27, 28, 29, 30], plays a crucial role in generating this co-occurrence.

## 2 Results

### 2.1 Self-organization of power law in PCA

To investigate the spontaneous emergence of power law in PCA, dissociated rat cortical neurons were cultured on high-density multi-electrode arrays (HD-MEAs) [20, 21], and their spontaneous activity was measured once a day (Fig. 1A). From the 26,400 recording electrodes in an area of 3.85 mm × 2.10 mm, up to 1,024 electrodes with the largest mean spike amplitudes of spontaneous activity were selected for 30-minute recordings. Spontaneous activity became detectable approximately 2 − 3 days in vitro (DIV). As reported in previous studies [26, 24, 31, 32, 33], bursts became observable as the culture matured (Fig. 1B).

**Figure 1:**
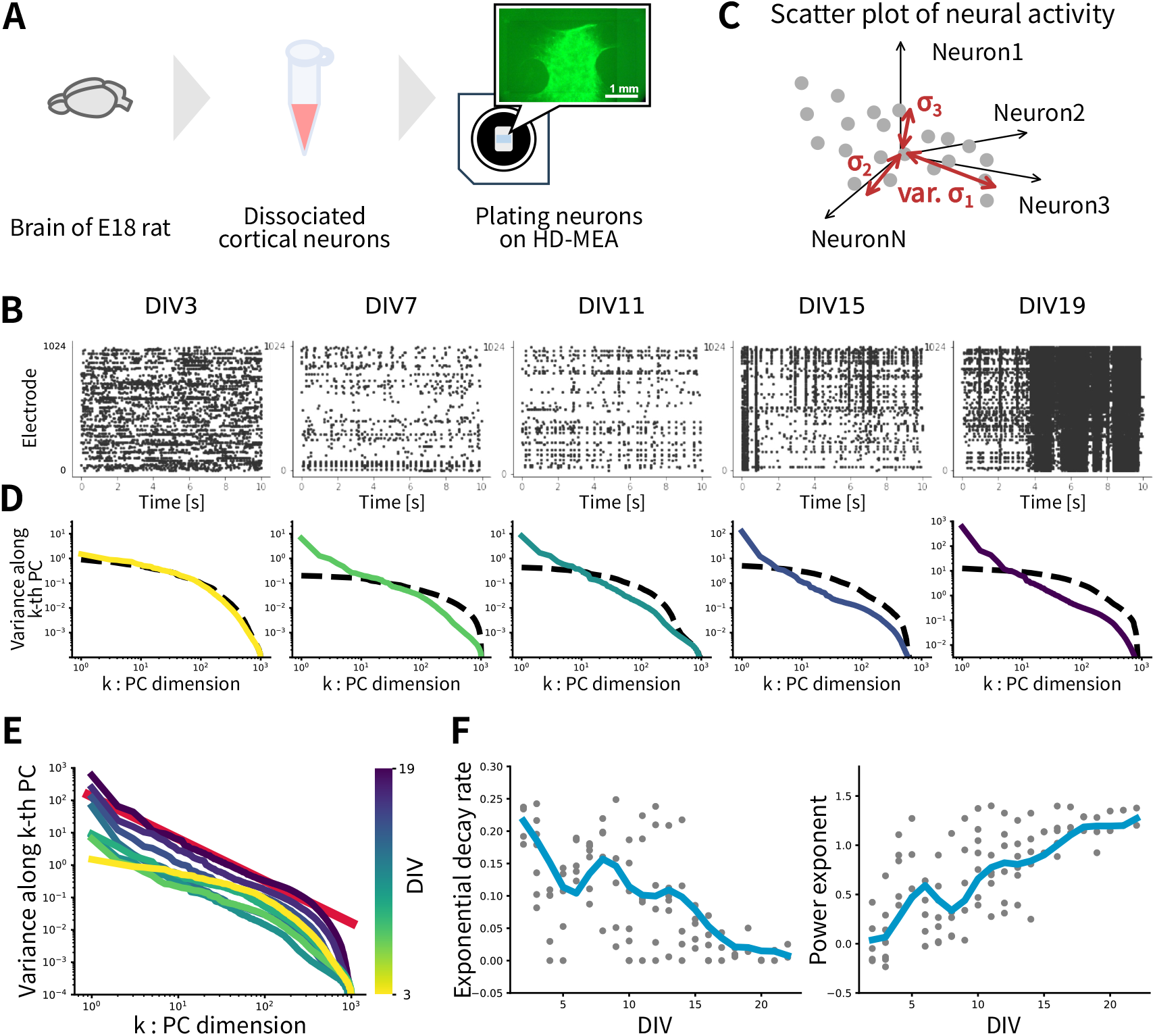
High-dimensional coordinated neural activity, characterized by a power-law decay of the explained variance in principal component analysis (PCA), is self-organized in the dissociated neuronal culture. (A) Dissociated neuronal culture. Brain tissue extracted from an E18 rat is dissociated, plated onto a HD-MEA, and cultured for several weeks. The inset shows a fluorescent image of GFP-labeled neurons on the recording electrodes. (B) Representative raster plots showing the evolution of spiking activities over DIV (days in vitro). (C) Outline of the analysis. The variance spectrum is obtained through PCA applied to a time series of high-throughput neural activity recorded from an HD-MEA for each dish on each day. Gray dots represent neural activity in the high-dimensional phase space at different time points of a dish on a day. (D) Variance spectra for each DIV. The vertical axis is the rank of the principal component (PC), i.e., the PC dimension. The horizontal axis is the variance along each dimension. Dashed lines are the same variance spectra obtained from shuffled data. The shuffling is performed as an independent circular shift of the time series for each electrode. (E) Superposition of the variance spectra for different DIV. The red line indicates a power-law fit, with an exponent of 1.30. (F) Exponential decay rate (left) and power exponent (right) derived from the variance spectra of each dish and each day. A lower exponential decay rate indicates that the decay of the variance spectra is well explained by a power law. Thick lines indicate 3-day moving averages of the data.

We performed PCA on 30 minutes of recorded neural activity and obtained variance spectra by ranking the variances along each PC in descending order (Fig. 1C). The analysis revealed that the power law in PCA was spontaneously established (Fig. 1D). Initially, the variance spectrum decayed more rapidly than a power function. However, as the culture matured, the spectrum began to follow a power function, appearing linear on a bilogarithmic scale. Figure 1E shows an overlay of these spectral transitions. These results indicate that, in this dish, the variance spectrum shifted from exponential decay to power-law decay around 10 DIV and remained stable at least until 19 DIV.

Averaging data across seven dishes revealed that power law in PCA was consistently established around 15 DIV. To quantify this transition, we fitted the spectra with the product of a power-law decay and an exponential decay (see Methods). The exponential decay rate, which quantifies the degree of exponential decay, was large in the early stages of development, indicating a dominance of exponential decay. As the culture matured, the decay rate decreased, reflecting a shift toward power-law decay (Fig. 1F). The power-law exponent stabilized at approximately 1.2. The emergence of the power law in PCA was consistently observed, independent of the temporal bin width used to vectorize the spiking activity (Fig. S1).

During this transition from exponential to power-law decay, spiking activity evolved from asynchronous firing to synchronous bursting (Fig. 1B). We hypothesized that the changes in variance spectra were driven by the emergence of synchronous firing. To test this hypothesis, we performed PCA on temporally shuffled neural activity. During the initial development period, shuffling did not alter the variance spectra. However, after the emergence of the power law in PCA, shuffling significantly disrupted the power law, reverting them to exponential decay (Fig. 1D). These findings indicate that the temporal structure of neural activity, including synchronized firing, is critical for the emergence of power law in PCA in the dissociated neuronal culture.

These findings indicate that the power law observed in PCA arises within dissociated neuronal cultures. Temporal correlations between electrodes driven by synchronized firing play a critical role in establishing this phenomenon. This suggests that the power law in PCA is a self-organized property of neural networks that emerges spontaneously during network formation.

### 2.2 Self-organization of spatial structure of PCs

A previous study of the spontaneous activity of the mouse cerebral cortex revealed that dominant PCs correspond to neural activity localized within specific brain regions, while minor PCs are associated with sparsely distributed, global neural activity (Fig. 2A; [7]). To investigate whether this characteristic spatial structure of PCs in high-dimensional population activity emerges spontaneously, we analyzed the spatial features of the data from the dissociated neuronal culture. The spatial distribution of each PC was obtained by mapping the weights of the PC with the spatial extent of electrodes.

**Figure 2:**
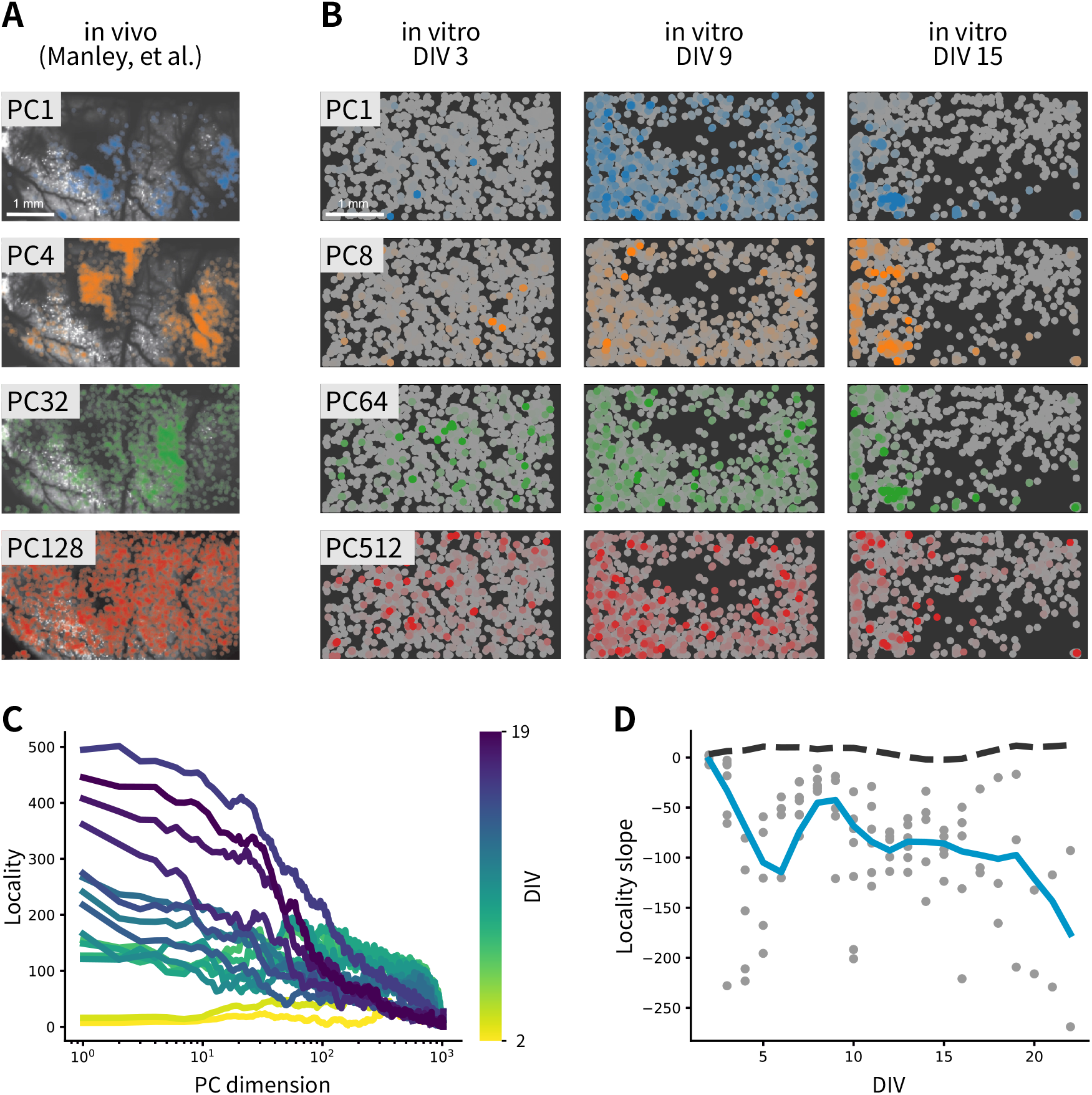
Self-organization of the characteristic spatial structure of principal components (PCs). (A) Spatial structure of PCs observed in a previous in vivo study [7]. Dominant, or low rank, PCs correspond to neural activity localized to specific brain regions, whereas minor, or high rank, PCs correspond to neural activity sparsely distributed across the cortex. (B) Spatial structures of PCs observed in the dissociated culture at early (DIV3), middle (DIV9), and late (DIV15) stages of the development. (C) Locality spectra of PCs at different DIVs, with colors representing the DIV. Thick lines indicate moving averages of the locality index with a window width of 10 PC dimensions. (D) Slope of the locality spectra (gray dots). The blue thick line shows a 3-day moving average of the slope. The dark gray dashed line indicates the locality slope for the data obtained from the temporal shuffle, which removes inter-electrode correlations.

Our analysis revealed that a characteristic spatial structure, similar to that observed in in vivo experiments, gradually emerged during development. In the early stages of development (3 DIV), all PCs corresponded to a small number of scattered electrodes. However, as the culture matured (15 DIV), dominant PCs began to correspond to localized clusters of electrodes, while minor PCs remained associated with globally distributed, sporadic electrodes (Fig. 2B).

To quantify the formation of this spatial structure, we employed “local Moran’s I”, a measure that evaluates the degree of spatial localization around a specific point in space. The spatial locality of PCs was assessed by calculating local Moran’s I for each electrode across all PCs and by counting the number of values exceeding a predefined threshold. Initially, the locality spectrum was flat, but as culture matured, it became steeper (Fig. 2C). This trend indicates the self-organization of the spatial structure of PCs, resembling that observed in vivo. Additionally, analysis of the slope of the locality spectrum confirmed the progressive emergence of spatial structure, which was absent during the initial stages of development (Fig. 2D). The formation of the spatial structure of the principal components was observed regardless of the choice of the two parameters used for the locality analysis (Fig. S2).

Similar to power law in PCA, the spatial structure of PCs was disrupted by temporal shuffling (Fig. 2D). Specifically, applying a circular shift to data with an established spatial structure, which removed inter-electrode correlations, disrupted the structure: the locality spectrum became nearly independent of the PC indices. These findings suggest that the spatial structure of PCs, like the power law in PCA, emerges from inter-electrode correlations driven by synchronous firing.

### 2.3 Co-occurrence of power law in PCA and neuronal avalanches

To further investigate the formation mechanism of the power law in PCA, we focused on neuronal avalanches, which represent a power-law distribution of synchronized firing event sizes (Fig. 3A). Neuronal avalanches are characterized by the power-law behavior of cascade size *S* and its probability *P* (*S*), following *P* (*S*) ∼ *S*^*-α*^. This phenomenon, which is self-organized in the brain and culture, has been widely studied as an example of SoC, as it emerges in a critical state when the excitatory/inhibitory balance is maintained. While the power law in PCA and neuronal avalanches are distinct phenomena that have been studied independently, we demonstrate below that these two phenomena co-occur. Based on this finding, we propose a possible mechanism underlying the emergence of the power law in PCA.

**Figure 3:**
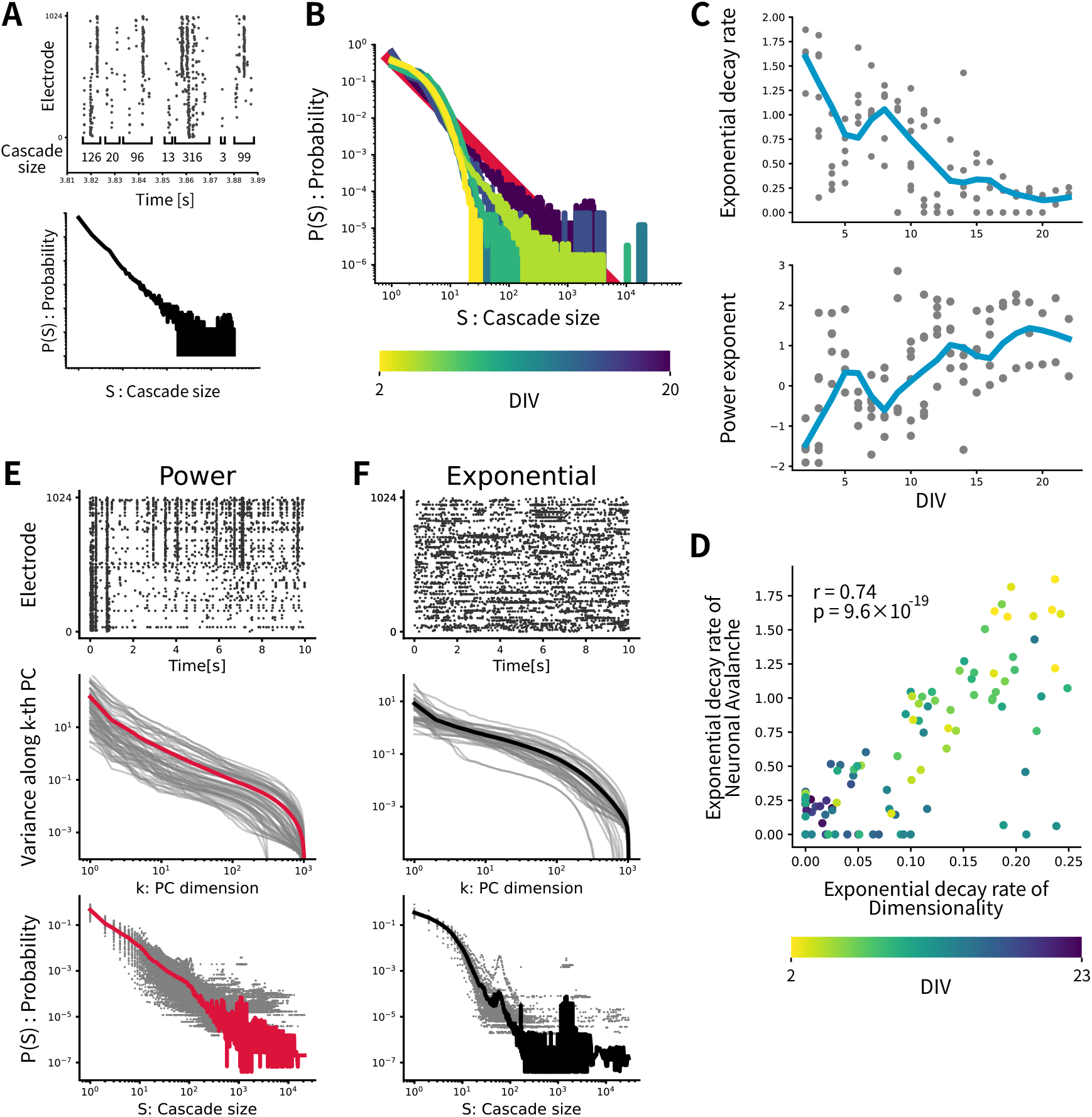
The power law in PCA is self-organized during the same period as neuronal avalanches in the dissociated neuronal culture. (A) Neuronal avalanches refer to the phenomenon where the distribution of cascade sizes follows a power law, where a cascade is defined as a sequence of consecutive neuronal firings. (B) Cascade size distributions for each DIV. (C) Exponential decay rate (top) and power exponent (bottom) of the cascade size distributions. Similar to Figure 1F, the lower the exponential decay rate, the better the decay of the distribution is explained by a power law. Thick lines represent 3-day moving averages of the data points. (D) Correlation between the exponential decay rate of the variance of the PCA and that of the neuronal avalanches. The horizontal and the vertical axes represent the exponential decay rate of the variance spectrum in the PCA and the cascade size distribution of the neuronal avalanches, respectively. (E,F) Representative raster plots (top), power-law in PCA (middle), and neuronal avalanches (bottom) for the “Power” group and the “Exponent group”, where data are classified into the two groups based on how well the power-law fits the variance spectrum in PCA.

We first confirmed the self-organization of neuronal avalanches in our dissociated neuronal culture. Figure 3B shows an example of the evolution of the cascade size distribution in a dish as development progressed. Data summarized from seven dishes show that the distribution shifted from an exponential decay to a power-law decay at approximately 15 DIV, with an exponent around 1.5 (Fig. 3C), consistent with previous studies [22, 23]. We confirmed that the self-organization of neuronal avalanches was robust to the temporal threshold for detecting cascades (Fig. S3).

We then compared the temporal evolution of the spontaneous organization of the neuronal avalanches with that of the power law in PCA. To quantify the evolution of the formation of these two power laws, we used the the exponential decay rate, which we used in Figure 1F to characterize the transition from exponential to power-law decay of the variances in PCA. Figure 3D shows the result. A clear positive correlation between the two exponential decay rates indicates that two power laws are established concurrently in the dissociated culture.

To further examine this co-occurrence of high-dimensional coordination and neuronal avalanches, we classified the recorded neural activity data from different DIVs and dishes into two groups based on their exponential decay rate of the variances in PCA. Recorded neural activity with exponential decay rate lower than a threshold value of 0.1 were classified into the “Power law” group, while all others with the higher values were classified into the “Exponential” group. Interestingly, although these groups were categorized based solely on the power law in PCA, regardless of the power law in the neuronal avalanches, it was clearly observed that both phenomena appeared in the Power-law group (Fig. 3E). In contrast, neither power law was observed in the Exponential group (Fig. 3F). A similar co-occurrence was also observed when data were classified based on the neuronal avalanches (Fig. S4). Together with the correlation shown in Figure 3D, these results provide strong evidence that the power law in the PCA and the neuronal avalanches emerge concurrently. This co-occurrence is notable because it is easily disrupted by temporal shuffling, reflecting that these phenomena are inherently unassociated (Fig. S5).

### 2.4 Heavy-tailed connectivity explains the co-occurrence

To investigate the mechanism underlying the co-occurrence of the power law in PCA and neuronal avalanches, conducted simulations and analyses using a mathematical model of a neural network (Fig. 4A).

**Figure 4:**
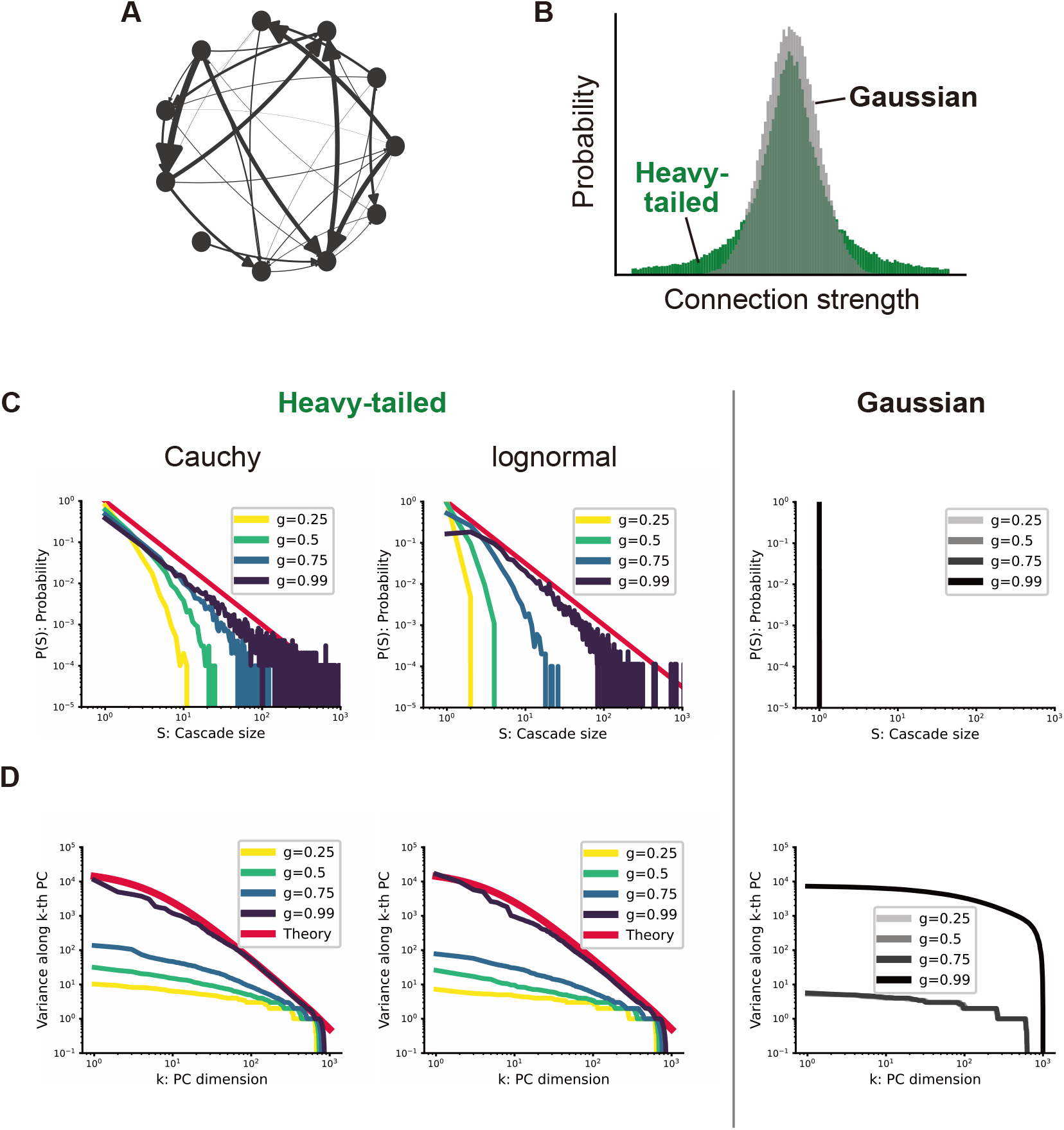
A network model with heavy-tailed connectivity explains the co-occurrence of the power-law in PCA and the neuronal avalanches. (A) Schematic of the recurrent neural network model. (B) Connections are sampled from a Gaussian distribution (gray) or a heavy-tailed Cauchy distribution (green). (C) Cascade size distributions for varying connection strengths: Cauchy distribution (left), lognormal distribution (center), and Gaussian distribution (right). The red lines represent the theoretical distribution. Only the networks with connections following the heavy-tailed distribution realize the power-law distribution of the cascade size at the critical point. See also [34]. (D) Variance spectra for varying connection strengths: Cauchy distribution (left), lognormal distribution (center), and Gaussian distribution (right). The red lines represent the theoretical solution (See Method). Similar to (C), only the heavy-tailed connectivity can achieve a power law in PCA. Note that the two power laws are realized at the same connection strength of the heavy-tailed network.

The model consists of a network of *N* neurons, each of which can take a value of 0 or 1. Neurons interact through a connection matrix *J*, and the state of a neuron becomes 1 at the next time step if the weighted sum of neural activities, determined by *J*, exceeds a threshold. The phase transition of this model has been examined in a previous study [34]. When connections are sampled from a Gaussian distribution, the model exhibits a discontinuous transition from an ordered phase to a chaotic phase as the mean connection strength increases. In contrast, if the connection strengths follow a biologically plausible heavy-tailed distribution, the chaos-order phase transition becomes continuous. This enables the network to achieve a stable critical state at the edge of chaos. The previous study demonstrated that neuronal avalanches occur in this critical state, whereas they are absent when the phase transition is discontinuous, as observed in networks with Gaussian-distributed connections (Fig. 4B, C).

Using this network model, we investigated whether power law in PCA emerges with heavy-tailed connection strength distribution and, if it is the case, whether it co-occurs with the neuronal avalanches. The numerically obtained neural activities of the model were analyzed in the same methods applied to the culture data.

Our results revealed that networks with heavy-tailed connections reproduced the co-occurrence of the power law in PCA and neuronal avalanches (Fig. 4D). The variance spectrum exhibited an exponential decay when the typical connection strength was sufficiently smaller than the critical point. As the connection strength increased, the spectrum transitioned to a power-law decay at the critical point. These changes in the variance spectrum resemble to the formation process of the power law in PCA observed in dissociated neuronal culture (Fig. 1D). In contrast, Gaussian networks, which fails to reproduce neuronal avalanches, were also unable to reproduce a power law in PCA at any connection strength.

### 2.5 Any heavy-tailed connectivity can reproduce the co-occurrence

Furthermore, we analytically demonstrated that both neuronal avalanches and the power law observed in PCA consistently emerge whenever the coupling strength follows a heavy-tailed distribution, not limited to the Cauchy distribution. By exploiting the heavy-tailed nature of the coupling strengths, the original model can be approximated with a model that has binary synaptic connections (see Method). For this binary coupling model, we derived the covariance matrix analytically. As a result, the variance spectrum, given by the eigenvalues of the covariance matrix, closely matched that of the original model and exhibited a power-law decay at the critical point of neuronal avalanches (*g* = 1) (Fig. 4C,D). The power-law exponent at the critical state was -1.5. Importantly, since different heavy-tailed coupling strength distributions converge to the same binary-reduced model and reproduce the co-occurrence, the coexistence of neuronal avalanche criticality and the power law in the binary model holds independently of the precise details of the coupling strength distribution.

These findings indicate that both the power law in PCA and neuronal avalanches emerge when synaptic connections follow a heavy-tailed distribution, whereas neither phenomenon occurs under Gaussian-distributed connections. Since it has been widely confirmed that synaptic connections are heavy-tailed in biological systems [27, 28, 29, 30], it is plausible that the coexistence of neuronal avalanches and the power law observed in PCA universally arises in biological systems as well. These results strongly suggest that the two power laws observed in vitro are self-organized through the maturation of biologically plausible heavy-tailed connections, which is thought to occur during early developmental stages in vivo. Since changes in network structure could not be directly measured in the present experimental setup, experimentally elucidating the relationship between synaptic connectivity and the emergence of the power law in PCA remains an important direction for future research.

## 3 Discussion

In this study, we demonstrated that two hallmark features of high-dimensional spontaneous activity observed in vivo, namely the power law in PCA and the characteristic spatial structure of PCs, are self-organized in dissociated neuronal culture. In the early stages of development, the variance spectrum exhibited an exponential decay rather than a power law, and all PCs were distributed on electrodes sparsely located on the dishes. However, by approximately 15 DIV, the variance spectrum transitioned to a power-law decay, and PCs acquired a localized spatial structure, indicating the self-organization of the specific high-dimensional structure of the coordinated population activity.

Furthermore, we discovered an unexpected link between the power law in PCA and neuronal avalanches, which have been studied independently. Neural activity with power law in PCA consistently exhibited neuronal avalanches, and vice versa. Since it was easy to construct temporal sequences that exhibit only the neuronal avalanches without the power law in PCA, this co-occurrence of two seemingly independent power laws is a nontrivial and unexpected observation.

To investigate underlying mechanisms of the co-occurrence, we employed a mathematical model of a recurrent neural network. Results of numerical simulation showed that this co-occurrence is reproduced when synaptic weight distribution follows the Cauchy distribution. Using analytical calculations, we demonstrated that any heavy-tailed network that is biologically plausible give rise to the co-occurrence.

We note the temporal structure of high-dimensional spontaneous activity observed in vivo [7]. This temporal structure is characterized by dominant PCs exhibiting longer time scales, while minor PCs exhibit shorter time scales. However, our study using the dissociated neuronal culture was unable to replicate such a temporal structure. This may be attributed to the relatively low spatial resolution of HD-MEA, where multiple cells contribute signals to a single electrode. Notably, the previous in vivo study also reported an absence of temporal structure under conditions of low spatial resolution. Applying calcium imaging to dissociated neuronal culture might resolve the spatial resolution issue, potentially enabling the investigation of the self-organization of the temporal structures.

The formation process of high-dimensional neural activity can be explained by the developmental process of neuronal avalanches, as these two phenomena co-occur in our experiment. Previous studies have reported that the formation of neuronal avalanches in dissociated cultures is governed by the “integration-then-fragmentation” process of their firing patterns [26, 35]. In this process, large-scale bursts initially emerge between 7 and 10 DIV due to the formation of excitatory synapses. These bursts are subsequently fragmented by the development of inhibitory synapses, resulting in a power-law distribution of diverse firing patterns. Notably, the development of the spatial structure of PCs observed in this study, which characterizes high-dimensional neural activity, can be understood through the “integration-then-fragmentation” process. The global firing patterns observed across all PCs at 9 DIV correspond to the occurrence of large-scale bursts, that is, the integration phase of the process, while the later emergence of dominant PCs associated with local firing patterns can be attributed to the fragmentation phase, which generates smaller-scale firing events. Elucidating the formation process of the power law in PCA, which is the other characteristic of high-dimensional neural activity, remains a challenge for future research.

Our results on neuronal culture also suggest the self-organization of high-dimensional neural activity in vivo. Several lines of evidence support this hypothesis: the heavy-tailed distribution of synaptic weights reported in vivo [27, 28, 29], the self-organization of neuronal avalanches achieved by excitation/inhibition (E/I) balance [36, 37, 38, 39], and the widespread presence of high-dimensional neural activity across various species and brain regions [4, 6, 7, 8, 9]. During development, E/I balance is thought to be established during critical periods, implying that the emergence of the power law in PCA and the spatial structure of PCs may occur within this developmental window.

Although we demonstrated the self-organization of the power law in PCA, this does not imply that the high-dimensional organization of neural activity in vivo is unrelated to learning. It is naturally expected that while a prototype of the high-dimensional structure is self-organized during the early stages of the animal’s development, it is later modulated by external stimuli, adjusting the power exponent accordingly through the experiences of animals. In fact, previous studies have observed power law in PCA on response to external stimuli as well as spontaneous activity [3, 40], discussing its role in cortical information encoding. Investigating the effects of long-term learning from the external world, in addition to short-term responses, could enhance our understanding of the relationship between high-dimensional spontaneous activity and neural computation.

Recent studies have revealed important roles of the power law in PCA for neural computation. The power law with a critical exponent has been shown to achieve optimal computational performance [3, 41, 42], and its scale-free property allows the brain to achieve unbounded encoding capacity relative to the number of its neurons [40]. Considering our findings that the power law is self-organized even in the absence of explicit sensory stimuli in animals, we speculate that these essential advantages for neural computation have been acquired and maintained throughout evolution by natural selection. The widespread observation of the power law in PCA across vertebrates, including zebrafish [8], may also supports this hypothesis, suggesting that it emerged before the divergence of vertebrates, potentially in conjunction with the development of the central nervous system. Elucidating the evolutionary mechanisms underlying this high-dimensional structure will be an important task for future studies.

## Methods

### Dissociated neuronal culture

This study was conducted in strict accordance with *Guiding Principles for the Care and Use of Animals in the Field of Physiological Science* published by the Japanese Physiological Society. The experimental protocol was approved by the Committee on the Ethics of Animal Experiments at the Graduate School of Information Science and Technology, the University of Tokyo (Permit Number: JA22-3).

The cell cultures were prepared on high-density multielectrode arrays (HD-MEAs; MaxOne, MaxWell Biosystems). Before seeding, the MEAs were treated with 0.07% polyethyleneimine (PEI; Sigma-Aldrich) and 0.02 mg/mL laminin (Sigma-Aldrich). Embryonic rat cortices were dissected from E18 rats and dissociated in 2 mL of 0.25% trypsin-ethylenediaminetetraacetic acid (Trypsin-EDTA, Life Technologies). Cells were then isolated by trituration, and approximately 380,000 cells were seeded onto the HD-MEA with 0.6 mL of plating medium, consisting of 450 mL of Neurobasal (Thermo Fisher Scientific), 50 mL of Horse serum (Cytiva), 1.25 mL of GlutaMAX (Thermo Fisher Scientific), and 10 mL of B-27 (Thermo Fisher Scientific). Twenty-four hours after seeding, half of the medium was replaced with the growth medium, composed of 450 mL of DMEM (Thermo Fisher Scientific), 50 mL of Horse serum (Cytiva), 1.25 mL of GlutaMAX (Thermo Fisher Scientific), and 5 mL of sodium pyruvate (100 mM, Thermo Fisher). Subsequently, half of the medium was exchanged twice a week. All experiments were conducted in an incubator at 36.5 ^*°*^C and 5% CO_2_.

### Recording of neural activity

Action potentials were recorded using the HD-MEA, which is equipped with 26,400 electrodes spanning area of 3.85 mm ×2.10 mm, allowing simultaneous recording from up to 1,024 electrodes out of them [20, 21]. Prior to recording data for analysis, 30 seconds of spontaneous activity was recorded for all the 26,400 electrodes. Based on these recordings, up to 1,024 electrodes with the highest mean spike amplitudes were selected for further recording. Spontaneous activity was then recorded from the selected electrodes for 30 minutes and used for subsequent analysis.

### Data processing and principal component analysis

In this study, to focus on data from cultures that showed successful growth, we analyzed spontaneous activity data with frequencies greater than 0.03 Hz from dishes where spontaneous activity was continuously observed for more than 10 days.

By recording 30 minutes of neural activity using 1024 electrodes of an HD-MEA and binning the data into 100 ms intervals to calculate firing rates, a data matrix of size 1024×18000 was obtained. PCA (Principal Component Analysis) was performed on this data matrix, and the eigenvalues corresponding to the explained variance of PCs are obtained. The eigenvalues of the covariance matrix of these vectors correspond to the variances along each PC. The variance spectrum was obtained by plotting the eigenvalues in descending order as a rank plot.

### Spatial structure of PCs

By associating the weights of each PC on each electrode with the spatial coordinates of the electrodes, we obtained the spatial distribution of the PCs. To evaluate the extent of spatial localization for each PC, we used the local Moran’s I statistic [43]. The local Moran’s I for the *k*-th PC around the *i*-th electrode is

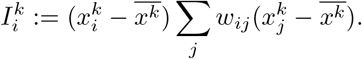

where 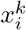 represents the weight of the *k*-th PC on the *i*-th electrode, and 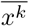denotes the average weight of the *k*-th PC across all electrodes. *w*_*ij*_ is the spatial correlation matrix of the electrodes, where pairs of electrodes separated by a distance *θ*_*r*_ = 200 µm or less are assigned a value of 1, and all others are assigned 0. The matrix is then row-normalized. To quantify the spatial locality of each PC, we calculated the number of electrodes for which the local Moran’s I exceeded a threshold value *θ*_*I*_ = 0.0001. We used this count instead of the global Moran’s I (the sum of all local Moran’s I values) to account for cases where the PC is localized around multiple distinct centers.

Furthermore, to evaluate whether there exists a spatial structure in which the dominant (minor) components exhibit local (global) distributions, we defined the locality slope as the slope of a linear approximation of locality spectrum plotted on a semi-logarithmic scale.

### Neuronal avalanches

To evaluate the formation of neuronal avalanches, characterized by a power-law distribution in cascade size, we defined cascades following methods from previous studies. For experimental data obtained from MEAs, a cascade was defined as a series of spikes with inter-spike intervals shorter than the mean inter-spike interval (mISI) [26]. For the mathematical model, a cascade was defined as the sequence of spikes triggered by a single neuron firing in the initial state [34].

### Parameter fitting

To evaluate the formation of power law in PCA and neuronal avalanches, we fitted each graph to a product of power-law decay and exponential decay:

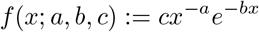

where *a, b, c* are optimized to minimize the mean squared error with respect to variance spectrum or cascade size distribution. This function asymptotically behaves as a power-law function for small *x* and transitions to an exponential function for large *x*. To prevent divergence at large *x*, the fitting was constrained to *b ≥*0. For neuronal avalanche, fitting was performed to cascade size 1 ≤ *S ≤*128.

The exponential decay rate *b* determines the range over which the power-law behavior is observed: when *b* is large, the range of power-law decay is narrow, while a value of *b* approaching zero corresponds to a broader range of power-law decay. Thus, we adopted the exponential decay rate *b* as a measure of power-law behavior.

### Mathematical model

Consider *N* neurons, each taking a binary value of 0 or 1. Neurons interact through a connection matrix *J*. If the weighted sum of inputs to the *i*th neuron at a time step 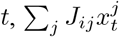, exceeds the threshold *θ*, the neuron’s state at the next step, 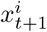, is set to 1:

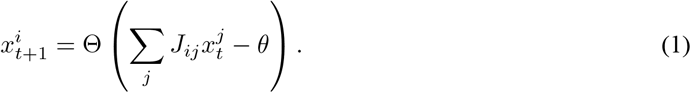

where Θ represents the Heaviside step function.

In this study, a connection probability was set to *p* = *O*(1/*N*) to achieve sparse connectivity. Cauchy, lognormal, and Gaussian distribution were employed as the probability distributions for non-zero connec-tion.

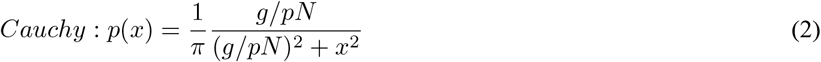

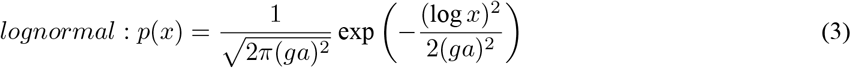

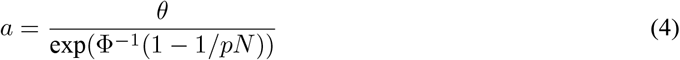

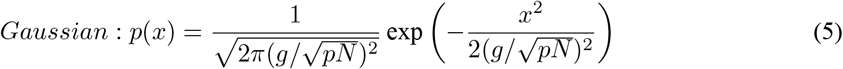

where *g* is the connection strength and Φ is the cumulative distribution function of the standard normal distribution. Coefficients are determined so that the phase transition occurs at *g* = 1. For details of phase transition of these models, refer to [34].

In numerical simulations with a finite number of neurons *N*, some network realizations exhibit abnormally high firing rates due to finite-size effects. Since this study focuses on the behavior in the large-*N* limit, we analyzed only those network realizations that did not show abnormally high firing rates in the simulations presented in Figure 4.

### Analytical Derivation of the Variance Spectrum

In this section, we derive the variance spectrum of a network model with heavy-tailed connectivity. The derivation proceeds in three steps: (1) approximating the heavy-tailed network by a network with binary connectivity, (2) evaluating the eigenvalue distribution of the covariance matrix of neural activities in the resulting binary network, and (3) deriving an analytical expression for the variance spectrum based on the eigenvalue distribution.

#### (1) Binary approximation of heavy-tailed networks

In networks with heavy-tailed connectivity, a small fraction of synapses are much stronger than the rest. In the low firing rate regime, simultaneous inputs from multiple presynaptic neurons are negligible, so postsynaptic firing is typically driven by these strong synapses. Therefore, in this regime, the heavy-tailed network can be approximated by a network with binary connectivity, in which synaptic weights are set to one if the corresponding synaptic strength in the original heavy-tailed network exceeds the firing threshold, and to zero otherwise (Fig. S6A):

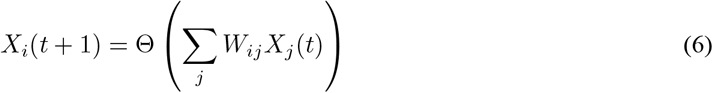

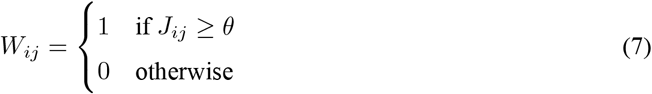

#### (2) Eigenvalue distribution of the covariance matrix of neural activity

The variance spectrum we consider is the rank plot of the variances of neuronal population activity along the principal component (PC) directions. These variances and PCs correspond to the eigenvalues and eigenvectors of the covariance matrix of neural activity, respectively. Thus, to derive the variance spectrum, we first evaluate the eigenvalue distribution, *P* (λ (*C*)), of the covariance matrix *C* of neural activity in the resulting binary network model (Fig. S6B).

In the low firing rate regime, the probability that two neurons fire simultaneously is low. Specifically, such co-firing typically occurs only when two post-synaptic neurons, *i*_1_ and *i*_2_, receive common inputs from a pre-synaptic neuron *j* (or via paths of length 2, 3, or more), a configuration that is statistically infrequent. Therefore, the off-diagonal elements of the covariance matrix are negligible compared with the diagonal elements, and the covariance matrix can be approximated by a diagonal matrix. Since the eigenvalues of a diagonal matrix are simply its diagonal entries, the eigenvalue distribution reduces to the distribution of the mean activity of neurons in the network, as follows:

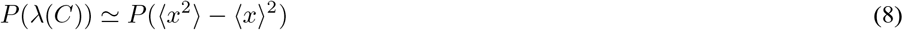

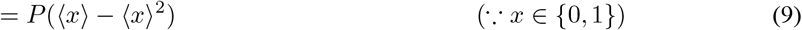

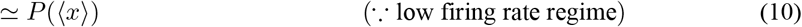

Interestingly, when the firing rate is low, the distribution of mean neuronal activity across the network is proportional to the tree-size distribution of a branching process defined on the network. To see this, note that neuronal activity is driven by rare and independent spontaneous firing events that trigger isolated cascades of activity. For a neuron to fire, a spontaneous event must occur either at the neuron itself or at one of its upstream ancestors, and the signal must successfully propagate to the neuron. Consequently, the mean activity of a neuron is proportional to the number of upstream nodes from which a cascade can reach that neuron. Therefore, up to a normalization constant, the distribution of mean neuronal activity across the network is proportional to the size distribution of upstream trees whose branches terminate at the neuron. We thus obtain

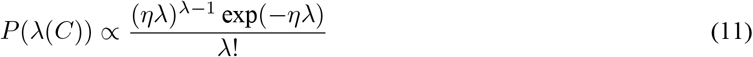

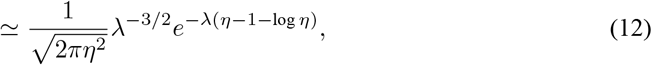

where the second line follows from Stirling’s approximation [44]. Here, ηdenotes the branching ratio, which is determined by the gain *g* of the original model and becomes η = 1 at the critical point *g* = 1.

#### (3) Variance Spectrum

We now derive the variance spectrum, i.e., the rank plot of the *N* eigenvalues drawn from the distribution *P*(*λ* (*C*)) and sorted in descending order (Fig. S6C). In the following analysis, we exploit the fact that λ takes discrete positive values and rescale it by its minimum value so that it can be treated as a positive integer.

Since Eq.(12) indicates that the eigenvalue distribution exhibits a transition from power-law decay with exponent -3/2 to rapid exponential decay at the crossover point *λ*\* = 1/(*η -1 -* log *η)*, we approximate the distribution by a truncated power law: *P*(*λ*(*C*)) ∼ *λ*^-3^/^2^ for *λ* < *λ*\*, and *P*(*λ*(*C*)) = 0 for *λ* > *λ*\*. Thus, the expected number of samples with *λ* = *k* is proportional to *k*^-3^/^2^, and if *n* denotes the number of samples with *λ* = 1, the expected number of samples with *λ* = *k* is *n/k*^*3/2*^*/* for 1 < *k* < *λ*\*.

In the rank plot, because the eigenvalues are sorted in descending order, the rank of an eigenvalue is given by the number of samples greater than or equal to it. Therefore, the *x*-coordinate corresponding to *λ* = *k* gives the expected number of samples greater than or equal to *k*, which is equal to the total number of samples *N* minus the number of eigenvalues strictly less than *k* (i.e., from 1 to *k* - 1). The rank plot is given in terms of *k* as:

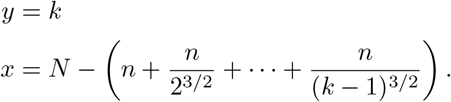

By eliminating *k*, we finally obtain the analytical expression of the variance spectrum

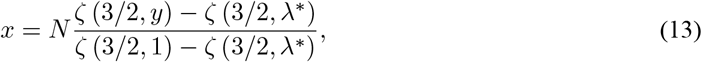

where 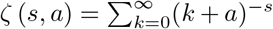 denotes the Hurwitz zeta function, and n = *N*/(*ζ* (3/2,1) - *ζ* (3/2, *λ*\*)) follows from the normalization condition.

The above derivation relies on the heavy-tailed nature of the connectivity distribution *p*(*J*), which guarantees the existence of rare but exceptionally strong synaptic connections. In the low firing-rate regime, these strong connections dominate the propagation of activity and justify the binary approximation introduced above. Since this mechanism depends only on the existence of such rare strong connections, the resulting variance spectrum is expected to be robust across a broad class of heavy-tailed connectivity distributions.

## Acknowledgments

This work is partly supportedby JSPSKAKENHI (24K15104,22K12186,23K21352,23H03465,24H01544, 24K20854), AMED (AMED-CREST JP23gm1510005h0003,24wm0625401h0001), the Asahi Glass Foundation, and the Secom Science and Technology Foundation.

## Declaration of generative AI and AI-assisted technologies in the writing process

During the preparation of this work the authors used ChatGPT in order to improve language and readability. After using this tool, the authors reviewed and edited the content as needed and take full responsibility for the content of the publication.

## Supplementary Figures

**Supplementary Figure 1:**
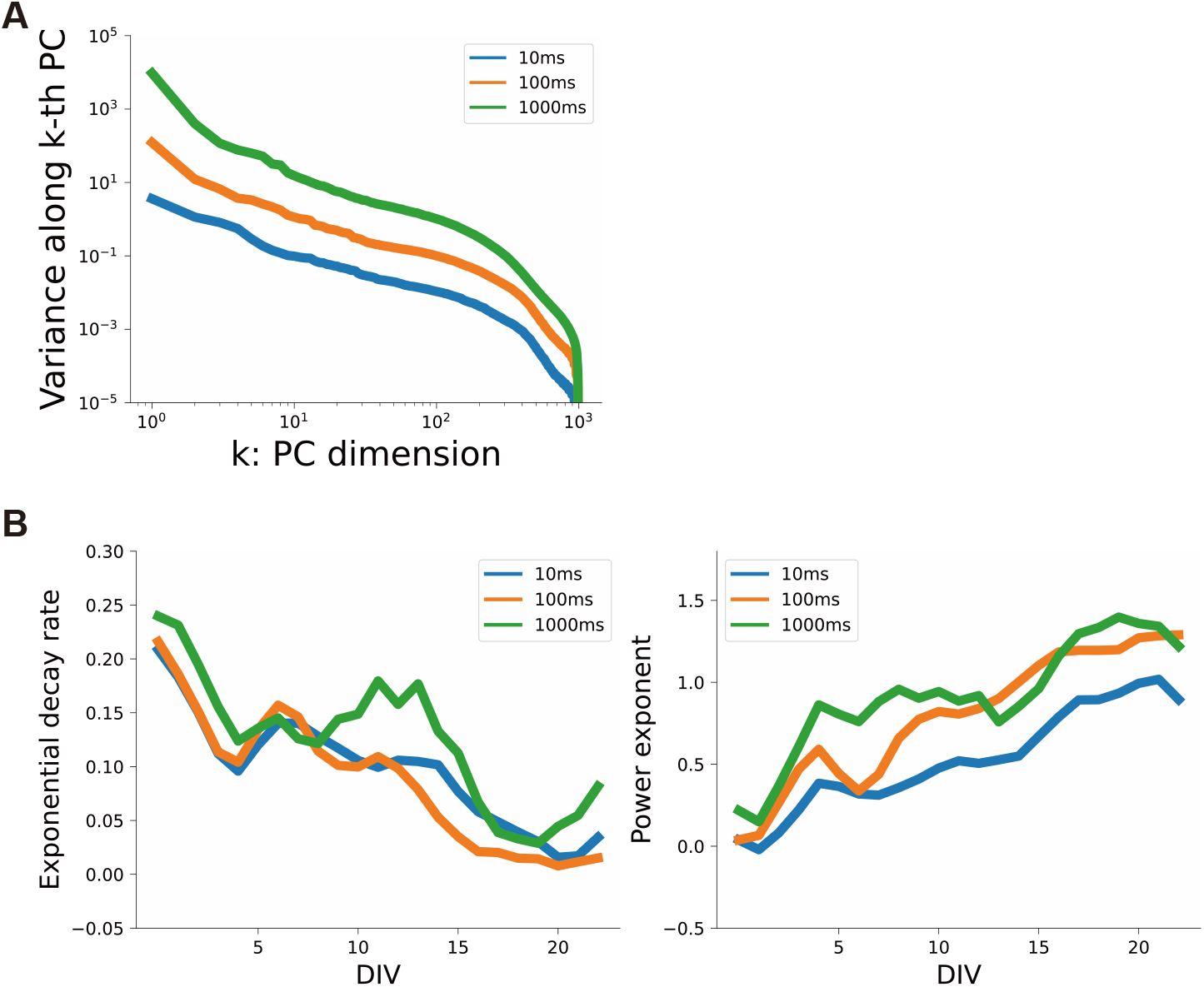
The emergence of power-law scaling in PCA is observed irrespective of parameter choices. (A) The power exponent of the variance spectrum remains unchanged even when varying the time bin width used to convert spike trains into firing rate vectors. (B) The qualitative behavior in which the variance spectrum transitions from exponential decay to power-law decay does not depend on the bin width. See also Figure 1B-C. We chose 100ms for other figures.

**Supplementary Figure 2:**
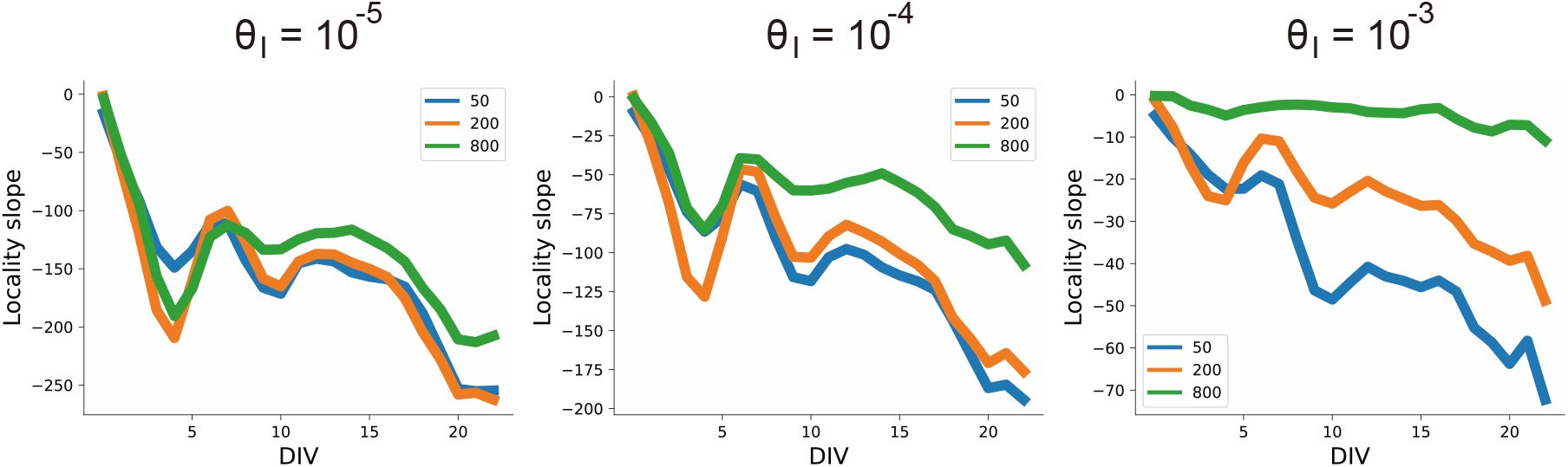
The emergence of the spatial structure of PCs is observed irrespective of parameter choices. The spatial structure remains unchanged even when varying the parameter *Q*_*r*_ and *θ*_***I***_ used to calculate locality. See also Figure 2D. We chose *θ*_*τ*_ *=* 200μm and *θ*_***I***_= 10^−4^ for other figures.

**Supplementary Figure 3:**
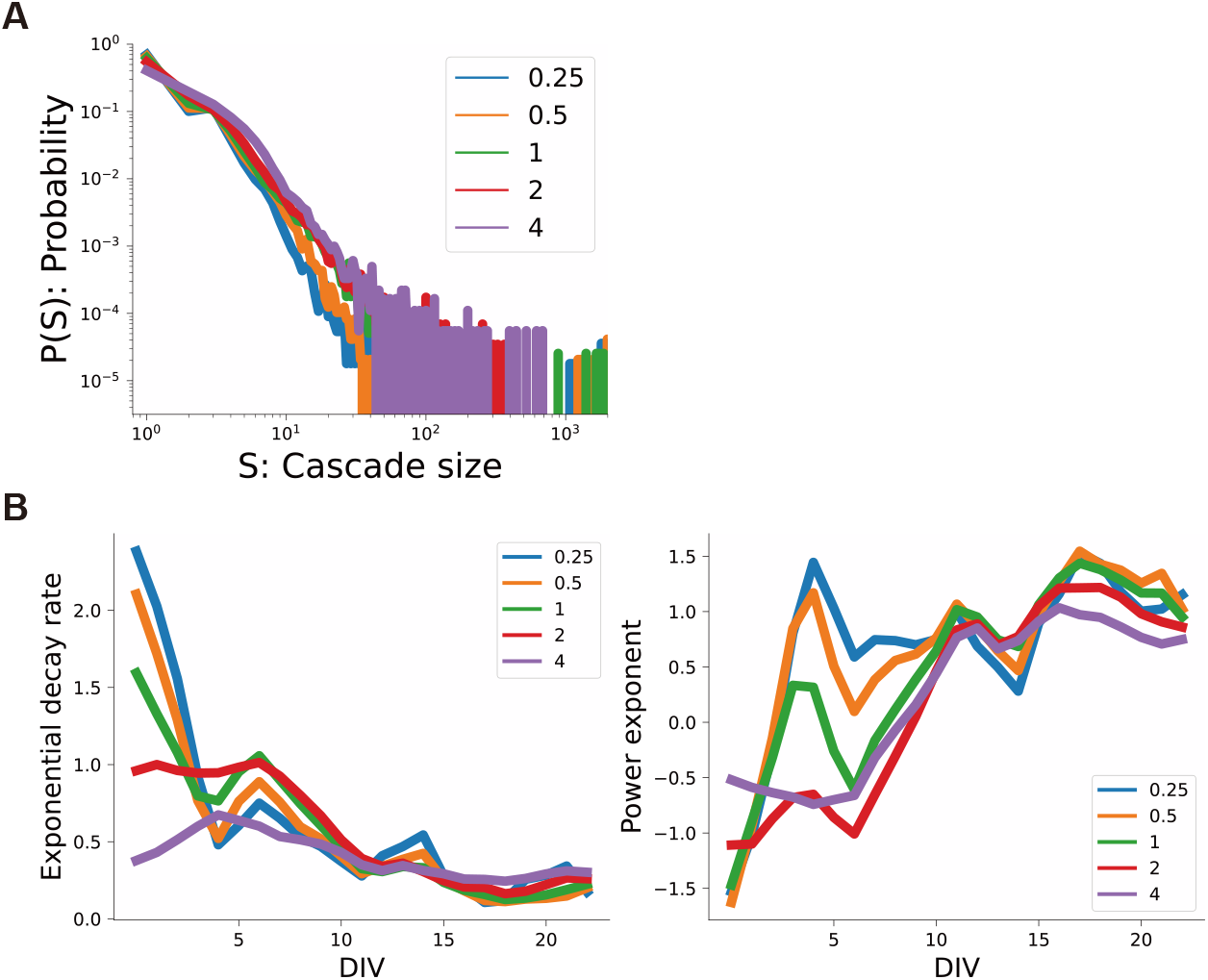
The emergence of neuronal avalanches is observed irrespective of parameter choices. (A) The power exponent of the variance spectrum remains unchanged even when varying the inter-spike interval threshold used to detect cascades. (B) The qualitative behavior in which the cascade size distribution transitions from exponential decay to power-law decay does not depend on the threshold. See also Figure 3C.

**Supplementary Figure 4:**
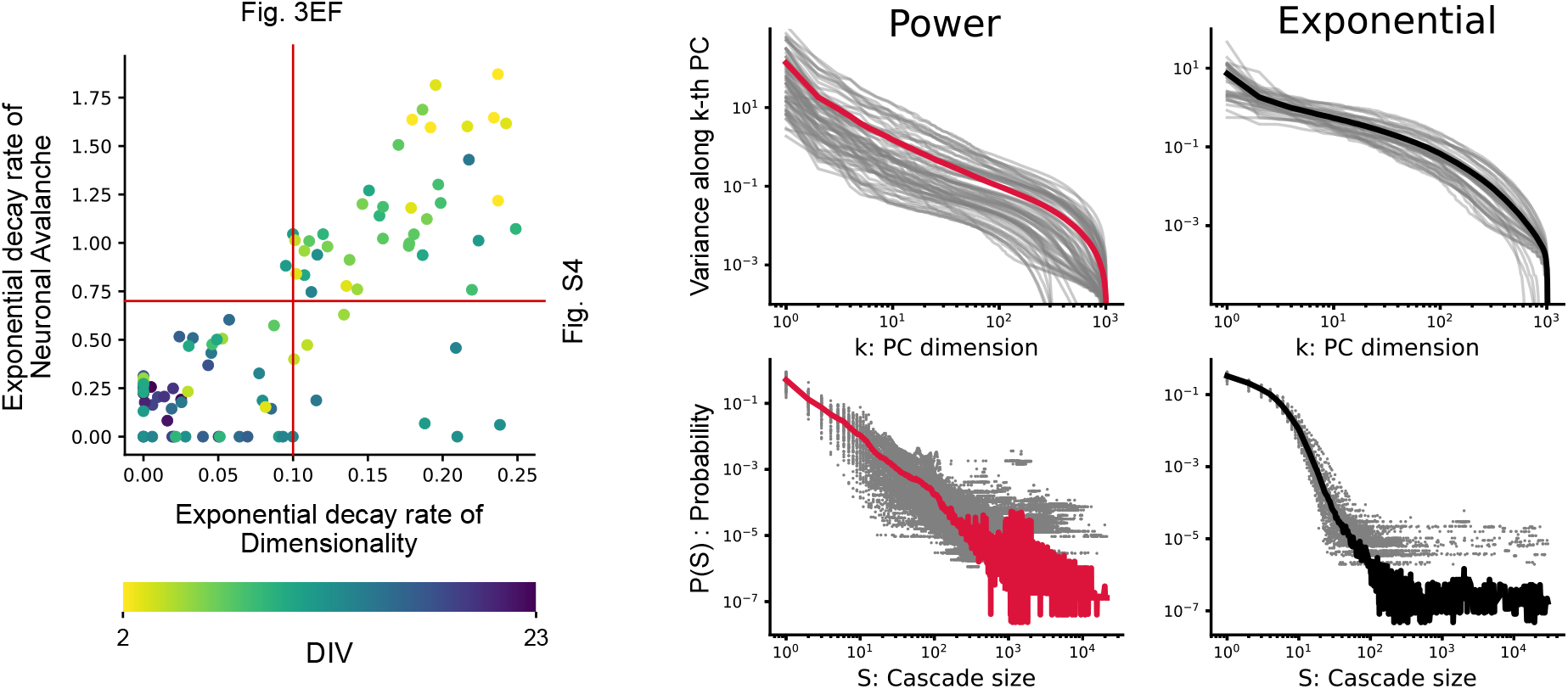
Co-occurrence of power-law dimensionality and neuronal avalanches is observed when data are classified based on the exponential decay rate of the cascade size distribution. See also Figure 3D-F.

**Supplementary Figure 5:**
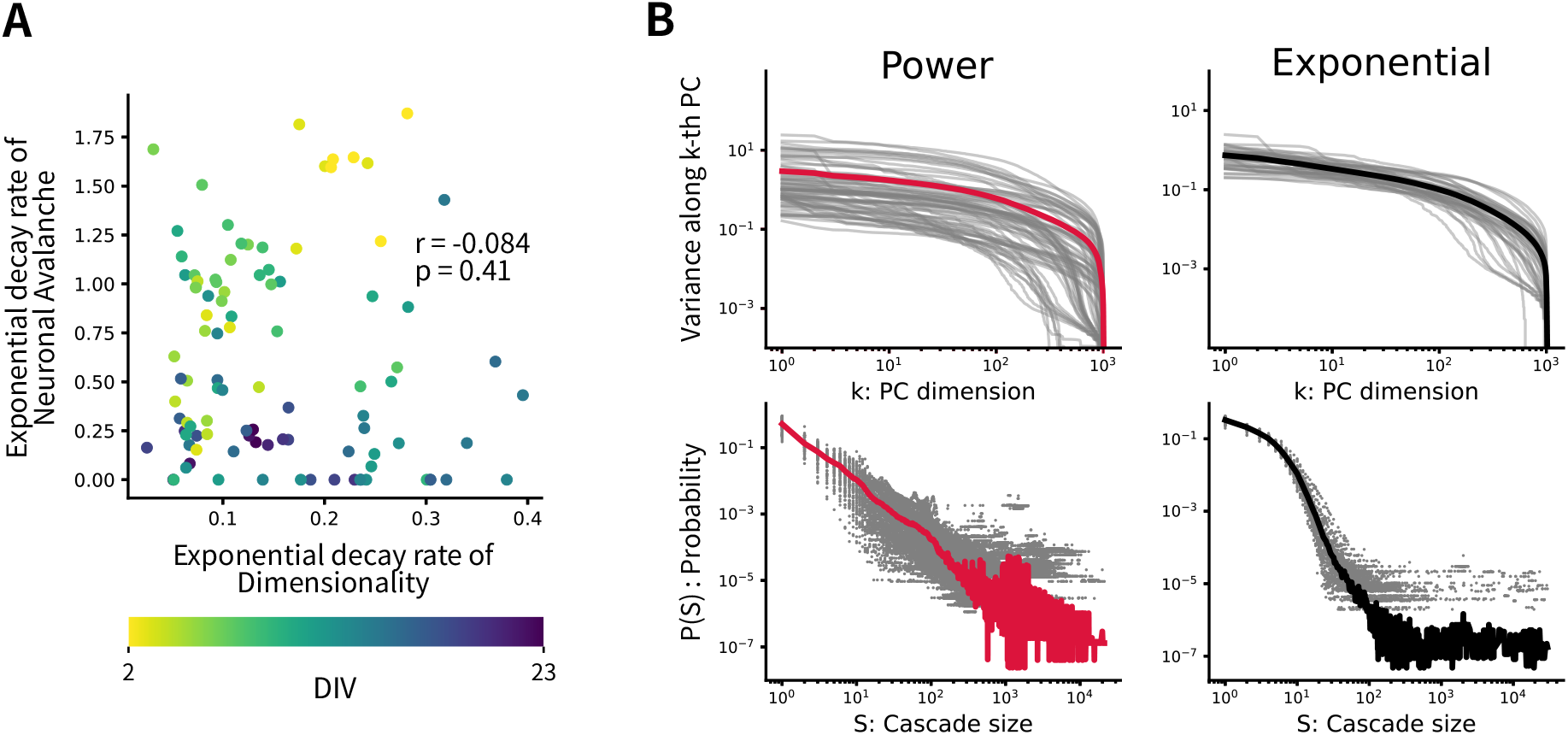
Co-occurrence of power-law dimensionality and neuronal avalanches is disrupted by shuffling on indices of electrodes. (A) Under shuffling, exponential decay rates show no significant correlation. (B)Shuffling disrupts the variance spectra, while the cascade size distributions remain unaffected.

**Supplementary Figure 6:**
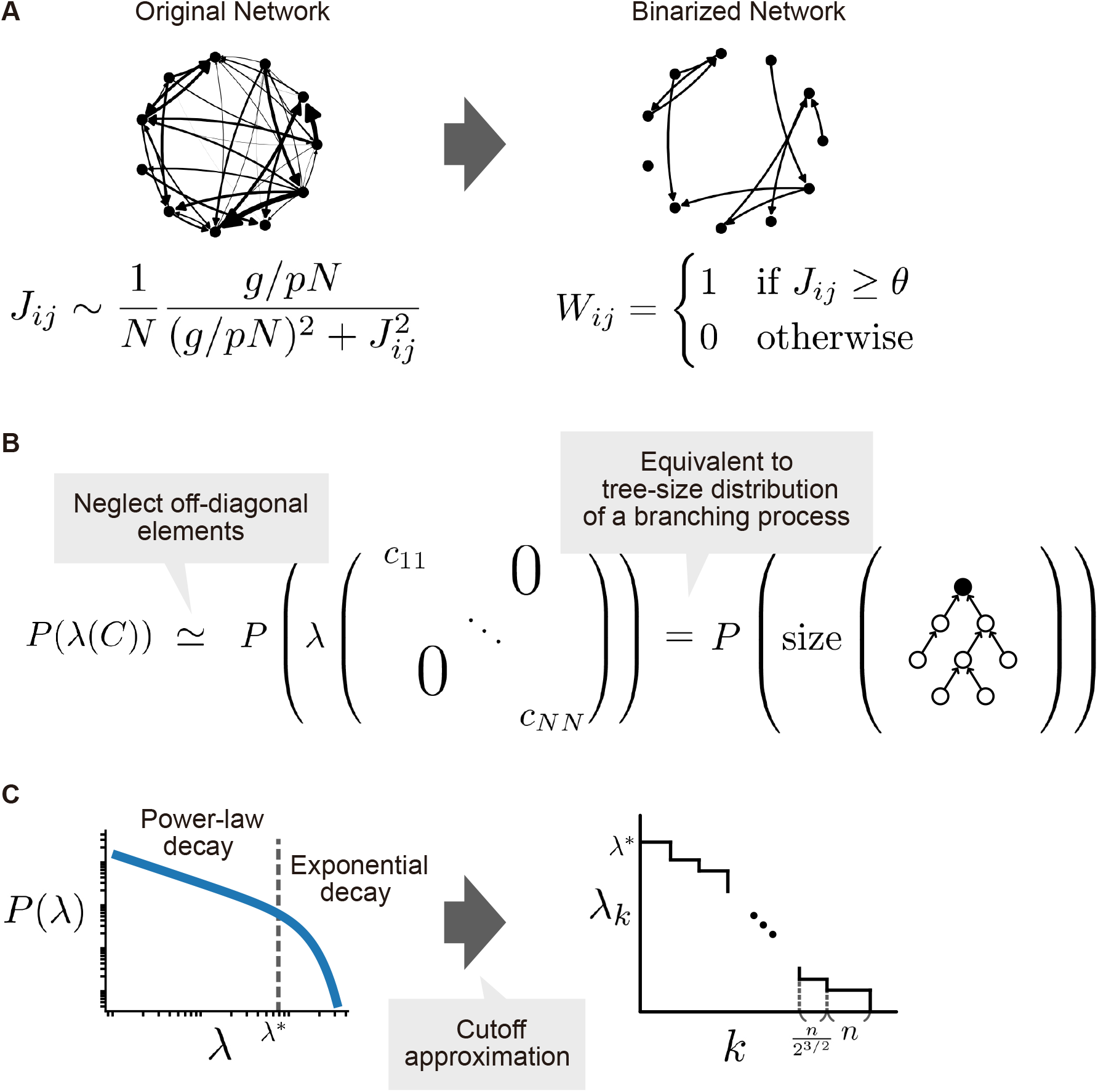
Steps for analytical derivation of the variance spectrum. (A) Binary approximation of the original network. (B) Eigenvalue distribution *P*(*λ* (*C*)) is equivalent to a tree-size distribution of a branching process. (C) Rank plot of the sorted eigenvalues *λ*_*k*_ can be derived using a cutoff approximation up to *λ*\*. See Methods for detailed derivations.

